# Metabolic transitions regulate global protein fatty acylation

**DOI:** 10.1101/2023.06.20.545712

**Authors:** Manasi Talwadekar, Subhash Khatri, Chinthapalli Balaji, Arnab Chakraborty, Nandini-Pal Basak, Siddhesh Kamat, Ullas Kolthur-Seetharam

**Author notes:** Corresponding authors: Siddhesh S. Kamat & Ullas Kolthur-Seetharam Email address. Equal contribution (Manasi Talwadekar, Subhash Khatri).

## Abstract

Intermediary metabolites and flux through various pathways have emerged as key determinants of post-translational modifications. Independently, dynamic fluctuations in their concentrations are known to drive cellular energetics in a bi-directional manner. Notably, intracellular fatty acid pools that drastically change during fed and fasted states act as precursors for both ATP production and fatty acylation of proteins. Protein fatty acylation is well regarded for its role in regulating structure and functions of diverse proteins, however the effect of intracellular concentrations of fatty acids on protein modification is less understood. In this regard, we unequivocally demonstrate that metabolic contexts, viz. fed and fasted states, dictate the extent of global fatty acylation. Moreover, we show that presence or absence of glucose, that influences cellular and mitochondrial uptake/utilization of fatty acids, affects palmitoylation and oleoylation, which is consistent with their intracellular abundance in fed and fasted states. Employing complementary approaches including click-chemistry, lipidomics and imaging, we show the top-down control of cellular metabolic state. Importantly, our results establish the crucial role of mitochondria and retrograde signaling components like SIRT4, AMPK and mTOR in orchestrating protein fatty acylation at a whole cell level. Specifically, pharmacogenetic perturbations that alter either mitochondrial functions and/or retrograde signaling affect protein fatty acylation. Besides illustrating the cross-talk between carbohydrate and lipid metabolism in mediating bulk post-translational modification, our findings also highlight the involvement of mitochondrial energetics.

## INTRODUCTION

Dietary inputs via metabolic intermediates have emerged as key drivers of cellular functions and hence impinge on physiological fitness. Specifically, the ability of primary and secondary metabolites to impact protein structure and function, across cellular compartments, is known to exert control over wide-ranging molecular mechanisms (1, 2). While protein fatty acylation is regarded as an abundant post-translational modification (3), metabolic contexts which dictate this globally are still unclear. Moreover, if/how *in situ* fluctuations in fatty acid concentrations contribute to protein acylation, for example, under fed and fast conditions, and its interplay with glucose metabolism and cellular energetics, is largely unknown.

Fatty acylation of proteins is known to regulate a multitude of processes. In addition to facilitating membrane association by increasing the hydrophobicity of a protein, fatty acylation of intrinsic membrane proteins affects structure, function, and their interactions (4). For instance, transient membrane interactions brought about by myristoylation (5–7) or palmitoylation (8–10) of proteins regulate key signaling processes in various cells. Further, fatty acylation of membrane-anchored receptors and transporters is known to mediate diverse cellular functions from neuronal synapses (11) to mitochondrial transporters and/or uncouplers (12–15). Not surprisingly, dysregulated fatty acylation has been implicated and is often thought to be central to the emergence of several diseases in humans (16–18).

It is interesting to note that unlike short-chain acyl moieties, which are largely formed by amide linkages of protein lysine residues (*N*-linked), long-chain fatty acyl chains seem to be conjugated either by amide or thioester bonds of protein lysine (*N*-linked) or cysteine residues (*S*-linked), respectively (19). However, there seems to be a bias for *S*-linked fatty acylation in the case of ≥ C16 fatty acid chains (20, 21), albeit this is not well established. Recent studies have demonstrated that acyl moieties can be covalently linked to proteins in an enzyme-dependent (22) and/or independent manner (23, 24). Enzyme-dependent fatty acylation has been largely attributed to *z*-DHHC palmitoyl acyl transferases (PATs) that catalyze the formation of a thioester bond between the thiol group of cysteine and fatty acids. Of the nearly twenty-three zDHHC PATs that have been categorized, most of them localize to ER and Golgi, and only a handful of them have been found to be present on the plasma membrane (25). Specific members of the DHHC family have also been shown to catalyze the addition of non-palmitate acyl-CoAs to cysteines of target proteins (26, 27). On the contrary, enzyme-independent fatty acylation of proteins involves a direct nucleophilic reaction between a cysteine thiol group and the thioester moiety of fatty acyl-CoA (28). It is important to note that, regardless of the mechanism, it is still not clear whether intracellular fatty acid concentrations can directly influence enzyme-dependent/independent protein acylation. Furthermore, given that fatty acylation is a reversible modification, relatively less is known about physiological contexts and mechanisms that mediate de-fatty-acylation. Earlier reports indicate a potential role for promiscuous thioesterases and NAD-dependent sirtuins in catalyzing the removal of *S*- and *N*-linked fatty acids from proteins. For example, while some α/β-hydrolase domain (ABHD) proteins have been recently reported to play a role in regulating protein *S*-acylation (29), SIRT6 is reported to possess a demyristoylase activity (30). Whether other members of these enzyme families play a role in de-fatty acylation remains unknown.

Independent of the mechanisms that mediate reversible fatty acylation, as mentioned earlier, intuitively, metabolic and physiological states are also likely to dictate the extent and nature of fatty acylation. Notably, various long-chain fatty acids such as palmitate, stearate, oleate, and arachidonate have been reported to reversibly modify proteins via cysteine linkages (31). This becomes interesting given that palmitate and oleate are the most abundant fatty acids in various cells and tissues in mammals, including humans (3, 32, 33). Moreover, in mammals, triglyceride composition, especially palmitate, stearate and oleate, have been shown to change dramatically under fed and fasted conditions in tissues such as adipocytes and muscles (34). Therefore, if/how physiological states lead to global changes in fatty acylation remains to be elucidated. Given this, our current study illustrates for the first time, a causal link between hepatic metabolism, energetics, and global protein fatty acylation. Our results describe the contribution of fed and fasted states and the involvement of mitochondrial functions in determining the extent of global protein fatty acylation. We provide unequivocal evidence, using pharmacological and genetic perturbations, that nutrient inputs exert direct control on fatty acid-dependent post-translational modifications of proteins.

## RESULTS

### Cellular metabolic status dictates protein fatty acylation

Although earlier studies have postulated a nutrient-dependent change in the fatty acylation of cellular proteins (35–39), whether this is governed by conditions that mimic fed or fasted states remain to be uncovered. Notably, glucose metabolism has also been shown to determine fatty acid flux in skeletal muscle (40). Therefore, we set out to investigate the impact of nutrient status on steady-state levels of protein fatty acylation across different cell types. We utilized three complementary approaches to assess fatty acylation viz. (a) exogenously administered clickable alkyne palmitate probe (C16-alkyne), (b) biotin exchange chemistry to label fatty acylated proteins, and (c) quantitative estimation of fatty acids released from proteins by liquid chromatography coupled to mass spectrometry (LC-MS/MS) (as detailed in the methods section) (**Figure 1**).

**Figure 1:**
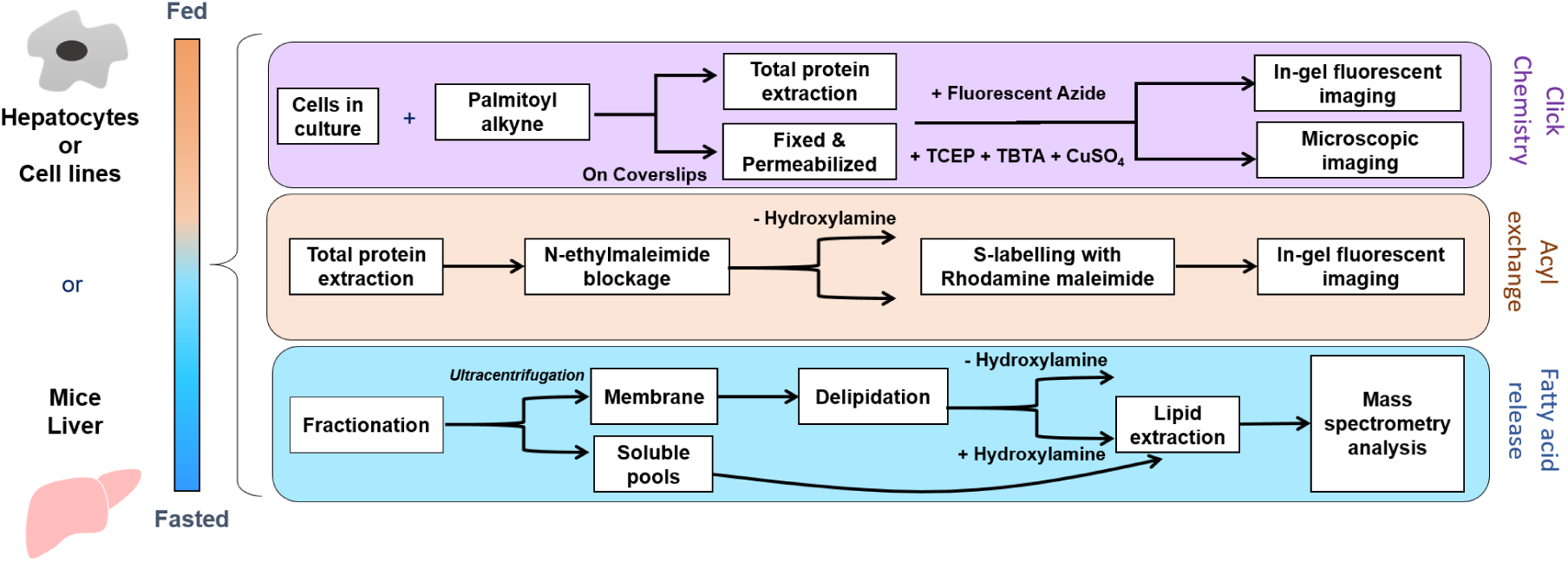
Methodology for assessing fatty acylation of proteins. Cells and mice tissues were subjected to fed and fasted paradigms and further processed using three methods: (i) Click chemistry or (ii) modified acyl exchange methods for qualitative assessment, as visualized by fluorescent gel imaging, and (iii) quantitative assessment for fatty acid release from proteins performed using LC-MS/MS analysis.

Differential nutritional states were mimicked by growing cells in media containing either low glucose (LG; 5 mM: Fasted state) or high glucose (HG; 25 mM: Fed State). To label fatty acylated proteins using click chemistry *in situ*, 100 µM C16-alkyne conjugated to BSA was exogenously added to the cells. Following this, protein lysates or fixed cells were fluorescently labelled using click chemistry by using a fluorescent azide probe (AlexaFluor 488). Enhanced protein palmitoylation (covalently linked C16-alkyne) was detected when cells were grown in low glucose conditions (**Figures 2A**, **2B**). Next, to assess the extent of endogenous fatty acylation, we employed hydroxylamine treatment for bulk biochemical measurements following acyl-exchange chemistry (**Figure 1**, **Supplementary Figure S1**) or quantitation of released fatty acids by LC-MS/MS (**Figure 1**). Hydroxylamine treatment has been employed to mostly report on *S*-linked fatty acylation (41). On assaying for hydroxylamine-sensitive fatty acylation levels, we found that the extent of global fatty acylation was significantly less under HG compared to LG conditions (**Figure 2C**). Since glucose/nutrient-dependent changes in fatty acid flux are both dynamic and pertinent in the liver, we quantified fatty acids released from proteins isolated from primary hepatocytes following hydroxylamine treatment. Consistent with earlier reports the cellular abundance of long chain fatty acids, namely palmitate (C16:0), stearate (C18:0), and oleate (C18:1) were higher than other fatty acid species, which were elevated in LG conditions (**Supplementary Figure S2**). More importantly, we found that hydroxylamine-sensitive fatty acylation of proteins was significantly enhanced under LG conditions with specific enrichment of palmitate (C16:0), stearate (C18:0), and oleate (C18:1) (**Figure 3A**), and these correlated well with corresponding increased cellular free fatty acid (FFA) pools (**Figure 3B**). To ascertain such nutritional state-dependent changes of global protein fatty acylation *in vivo*, we harvested liver from *ad libitum* fed or 24 hours fasted mice, and repeated hydroxylamine sensitive lipidomics analysis. Akin to results so far, both the protein fatty acylation (**Figure 3C**) and cellular FFA pools (**Figure 3D**) were significantly elevated in the fasted liver samples. Together, these findings clearly indicated a strong correlation between cellular metabolic status, such as fed and fasted state, and the extent of protein fatty acylation. More importantly, our data showed concordance between intracellular FFA pools and the enrichment of specific species of fatty acids, which were covalently linked to proteins.

**Figure 2:**
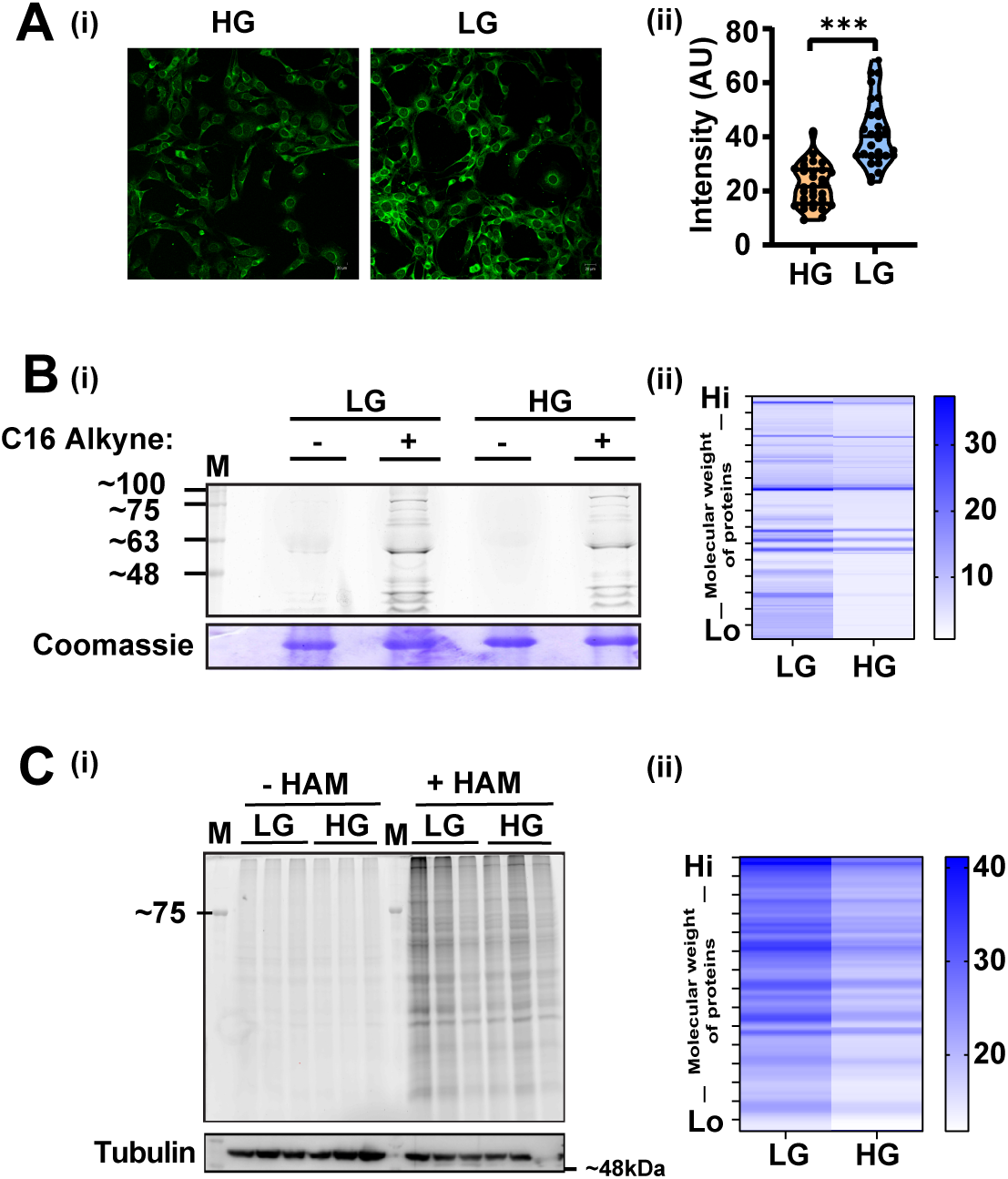
Nutritional states drive differential protein fatty acylation. **(A)** NIH 3T3 cells were grown under high glucose (HG) and low glucose (LG) conditions in the presence of C16-alkynyl fatty acid (FA) for 4 hours. Cells were fixed, permeabilized, and subjected to click chemistry reaction with AlexaFluor 488 azide: (i) Imaging done using confocal microscopy (scale bar 20 microns), (ii) Quantification of fluorescence intensity (n = 30). Data is analyzed by the Student’s t-test. A value of *p* ≤ 0.05 was considered statistically significant, \*\*\**p* ≤ 0.001. **(B)** HEK293 cells were treated with 100 µM C16-alkynyl FA probes under LG and HG conditions. Controls were treated with DMSO instead of FA. Cellular proteins were extracted and reacted with AlexaFluor 488 azide (N = 2): (i) Samples were resolved using gel electrophoresis and in-gel fluorescence was visualized using a Typhoon imaging system. (ii) Heat map depicting lane quantitation of band intensities. **(C)** Acyl exchange done using protein lysates (N=2, n=3): (i) C2C12 myoblasts grown in LG and HG media subjected to acyl exchange for total *S*-acylation using Rhodamine maleimide, tubulin used as loading control (ii) Heat map showing the mean of lane quantitation of band intensities for differential *S*-acylation in C2C12 cells in LG and HG.

**Figure 3:**
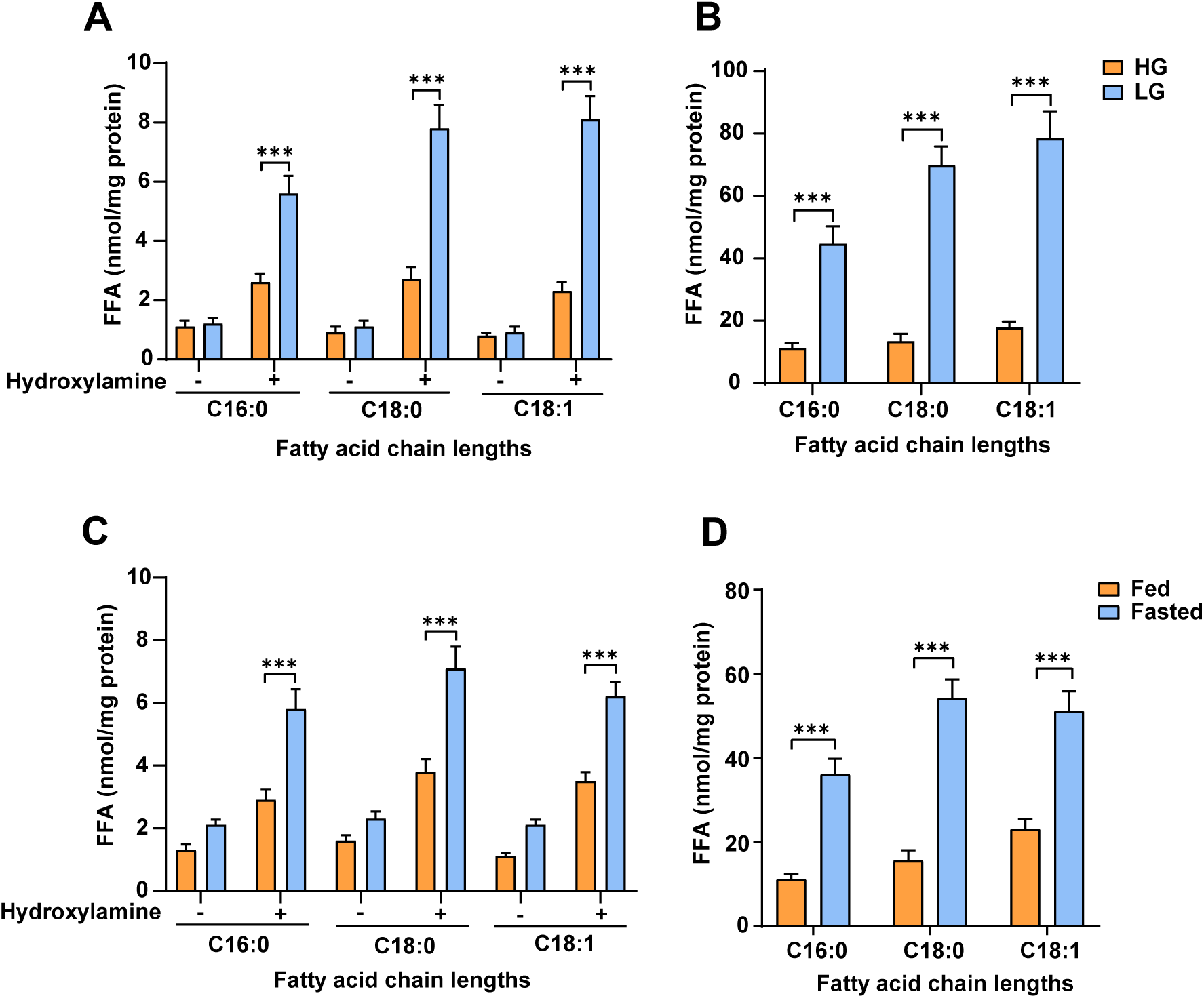
Estimation of protein fatty acylation in fed and fasted states in cells and tissues. **(A and B)** Primary hepatocytes were grown under HG and LG conditions (N=3, n=3) (**A**) Free fatty acid (FFA) released from membrane fractions after hydroxylamine treatment and (**B**) the associated cellular FFA pools. **(C and D)** Fatty acid content from liver tissue collected from 10 –12 weeks old *ad libitum* fed and 24 hours fasted C57BL6 mice (N=2, n=3) (**C**) post hydroxylamine treatment in membrane isolates and (D) their respective cellular FFA pools. Data are represented as mean ± SEM and analyzed by the Student’s t-test and ANOVA. A value of *p* ≤ 0.05 was considered statistically significant. \*\*\**p* ≤ 0.001.

### Protein fatty acylation is associated with fatty acid concentration-dependent uptake and utilization

Substrate availability has been shown to affect metabolite-derived protein modifications, as in the case of acetylation (42). In this context, we treated primary hepatocytes with increasing amounts of BSA-conjugated palmitate, which revealed a concentration-dependent change in intracellular levels of palmitate (**Supplementary Figure S3A**). We used this paradigm to assess the contribution of exogenously provided fatty acid on global protein acylation. Increased intracellular palmitate levels by the addition of 250 μM BSA-palmitate (**Figure 4A**), resulted in hydroxylamine-sensitive palmitoylation of proteins (**Figure 4B**). Further, to validate the dependence on fatty acid uptake, we treated primary hepatocytes with sulfo-N-succinimidyl oleate sodium (SSO), a CD36 inhibitor. CD36 is a key fatty acid transporter and has earlier been established to dictate intracellular FFA concentrations (43). SSO treatment of primary hepatocytes attenuated the increase in cellular palmitate levels caused by exogenously provided BSA-palmitate, indicating the inhibition of CD36 (**Figure 4C)**. More interestingly, consistent with our hypothesis, we found a significant reduction in palmitate-induced-protein palmitoylation following pharmacological inhibition of CD36 (**Figure 4D**). Moreover, the exogenously added palmitate did not lead to any effects on the cellular oleate, stearate, and extent of protein oleolylation or stearoylation, in our paradigm (**Supplementary Figure S3B, S3C, S3D).** In addition to providing strong corroborative evidence vis-à-vis substrate dependence, it also negated the contribution of lipogenic or fatty acid chain length elongation/desaturation mechanisms in driving global protein fatty acylation (**Figure 4E**).

**Figure 4:**
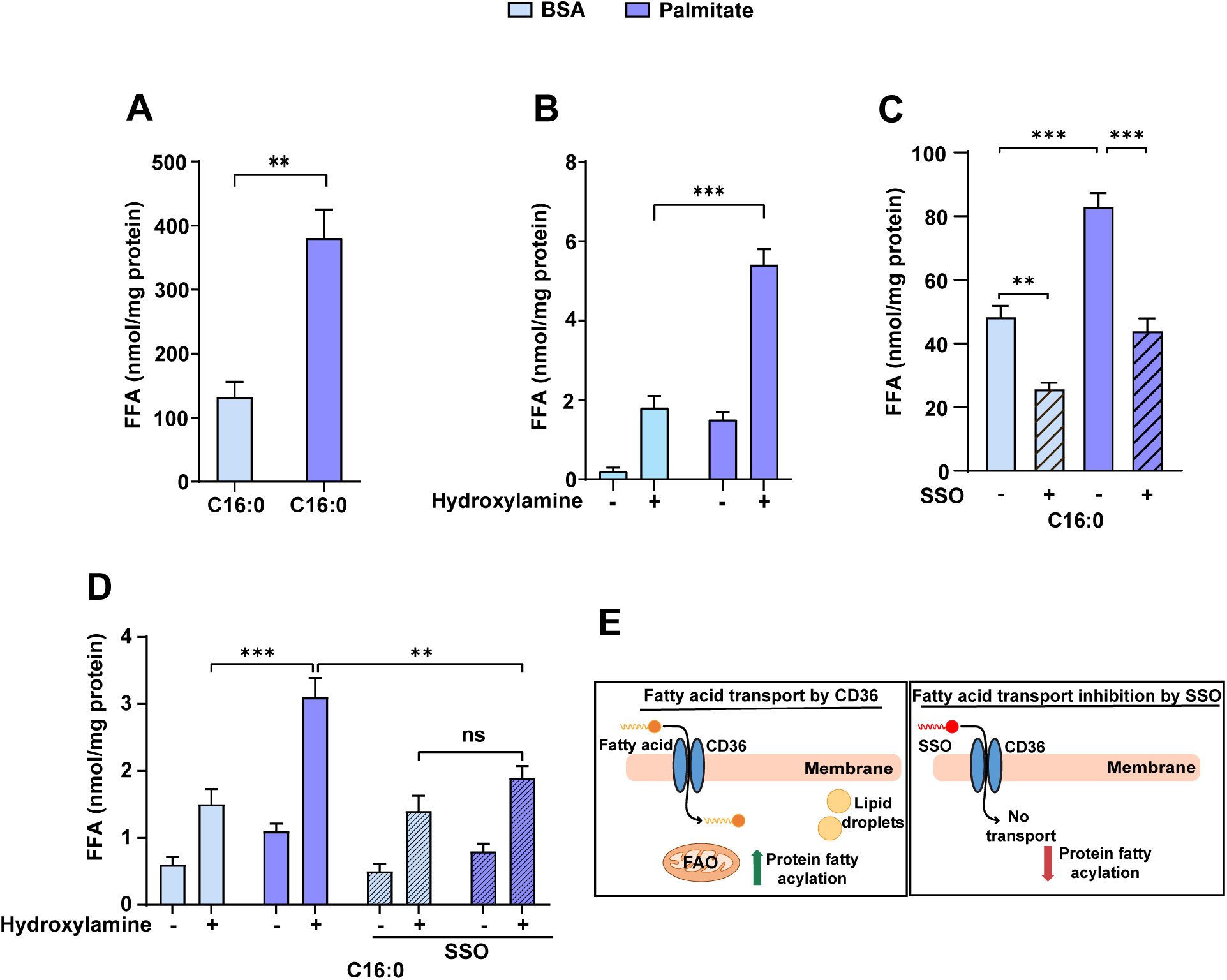
*S*-Fatty acylation of proteins is governed by substrate pools in a metabolic state. **(A and B)** Primary hepatocytes were grown in the presence of either BSA alone or 250 µM BSA conjugated palmitate (N=3, n=3). (**A**) Palmitate pools in the soluble fraction (**B**) Corresponding palmitate release assay for membrane-bound fatty acids **(C and D)** Primary hepatocytes grown in the presence of BSA conjugated palmitate in the presence and absence of CD36 inhibitor (SSO) (N=3, n=3) (**C**) Cellular palmitate levels and (**D**) Protein palmitoylation detected in membrane fraction after hydroxylamine treatment. **(E)** Depiction of cells treated with fatty acid transporter CD36 inhibitor, SSO. Data are represented as mean ± SEM and analyzed by the Student’s t-test and ANOVA. A value of *p* ≤ 0.05 was considered statistically significant. \*\**p* ≤ 0.01; \*\*\**p* ≤ 0.001.

As mentioned earlier, the acylation of proteins is proposed to occur both in an enzyme-dependent and independent manner with a paucity of mechanistic insights for the latter. In the case of long-chain fatty acylation, enzyme-dependent modification has been largely attributed to the activity of PATs. Thus, we employed 2-bromopalmitate (2-BP), a general inhibitor for PATs (**Figure 5A**). Surprisingly, 2-BP treatment had no impact on the extent of protein fatty acylation as assayed by acyl exchange chemistry (**Figure 5B**). Further, quantitative analyses of hydroxylamine-sensitive released fatty acids corroborated these findings (**Figure 5C** and **Supplementary Figure S4A**). Interestingly, the 2-BP treatment also did not affect the soluble pools of C16:0 (**Figure 5D**), C18:0, and C18:1 (**Supplementary Figure S4B**). Although these results suggest a predominant role of non-enzymatic covalent addition of fatty acids, this could be context-specific (hepatocytes grown in LG medium) and does not exclude the possibility of PAT-dependence in other physiological conditions. Together, results presented thus far suggest that the nutritional state drives intracellular concentration-dependent, non-enzymatic fatty acylation of proteins.

**Figure 5:**
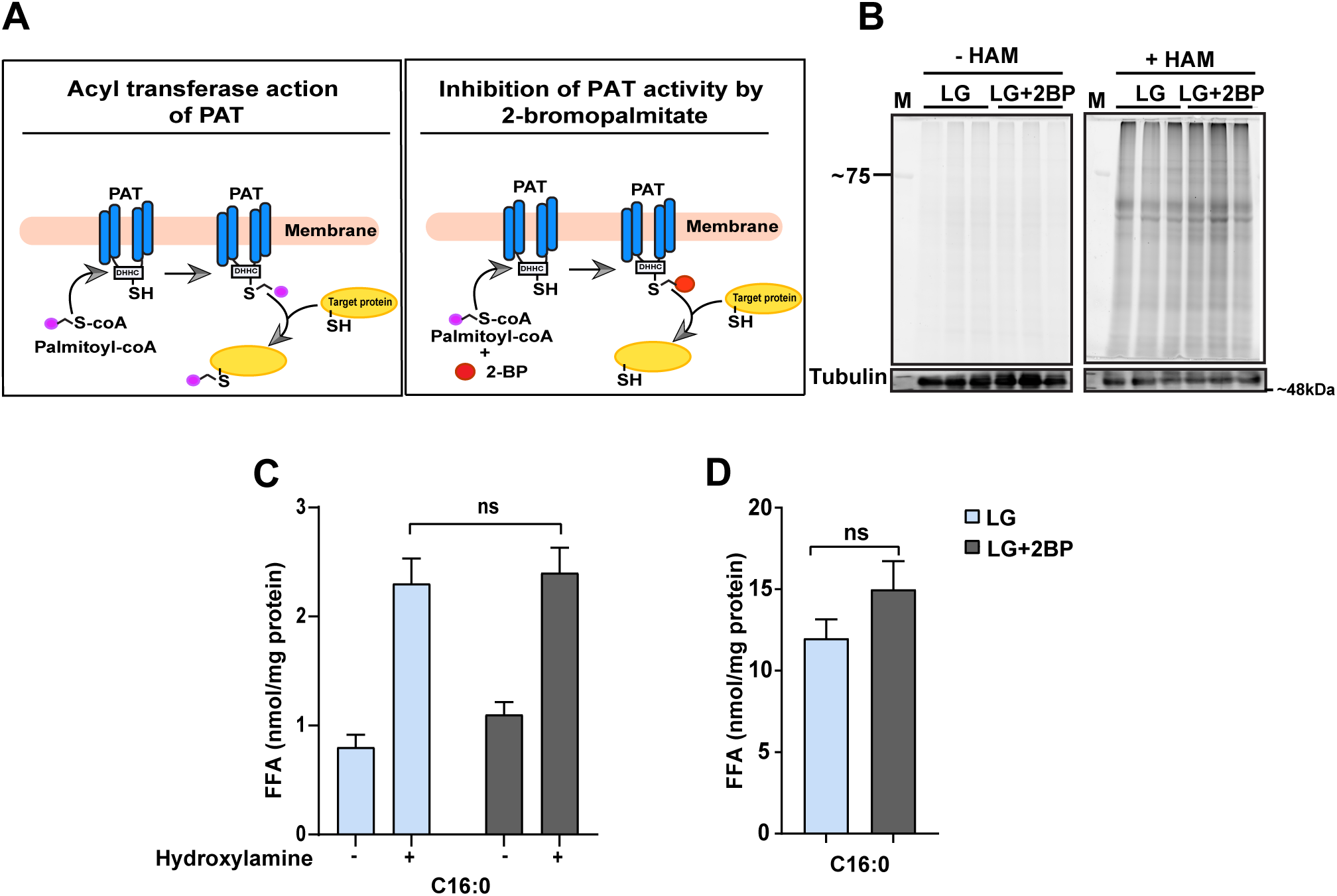
Contribution of non-enzymatic fatty acylation in the fasted state. **(A)** Schematic representation of the action of 2-BP on protein fatty acylation. **(B-D)** Primary hepatocytes were grown in LG conditions in the presence and absence of PAT inhibitor, 2-bromopalmitate (2BP): **(B)** Gel image showing total fatty acylation before and after hydroxylamine treatment (N=2, n=3) **(C)** Estimate of fatty acid released from proteins from the membrane fraction along with (N=2, n=3) **(D)** their respective cellular pools post 2-BP treatment (N=2, n=3). Data are represented as mean ± SEM and analyzed by the Student’s t-test and ANOVA. A value of *p* ≥ 0.05 was considered statistically non-significant (ns).

### Metabolic sensors that promote fasted state contribute to protein fatty acylation

Since nutrient availability seemed to dictate the extent of protein fatty acylation, we next investigated the involvement of important metabolic sensors viz. AMPK and TOR. To reiterate, because global acylation was higher in a fasting state, we employed chemogenetic perturbations to mimic a nutrient-deprived state as reported earlier (44, 45). As expected, rapamycin-mediated inhibition of mTORC1 (**Supplementary Figure S5A**) resulted in increased soluble FFAs (**Figure 6A**) and importantly, also the HAM sensitive release of FFAs from proteins (**Figure 6B**). This was despite hepatocytes being grown in HG medium, which otherwise showed lower protein fatty acylation (**Figure 3A**). Conversely, under LG conditions AMPK inhibition using compound C (**Supplementary Figure S5B**) resulted in a robust decrease in FFA levels both cellular (**Figure 6C**) and that released from proteins (**Figure 6D**). Here again, fatty acylation following AMPK inhibition that is known to counter the fasting response was comparable to that observed in cells grown in HG medium. To confirm this, we also activated AMPK by treating hepatocytes with AICAR (**Supplementary Figure S6A**) and FCCP (**Supplementary Figure S6B**) and found the opposite effect i.e., an increase in protein fatty acylation. Together, these results suggest an interplay between cellular energetics and global fatty acylation.

**Figure 6:**
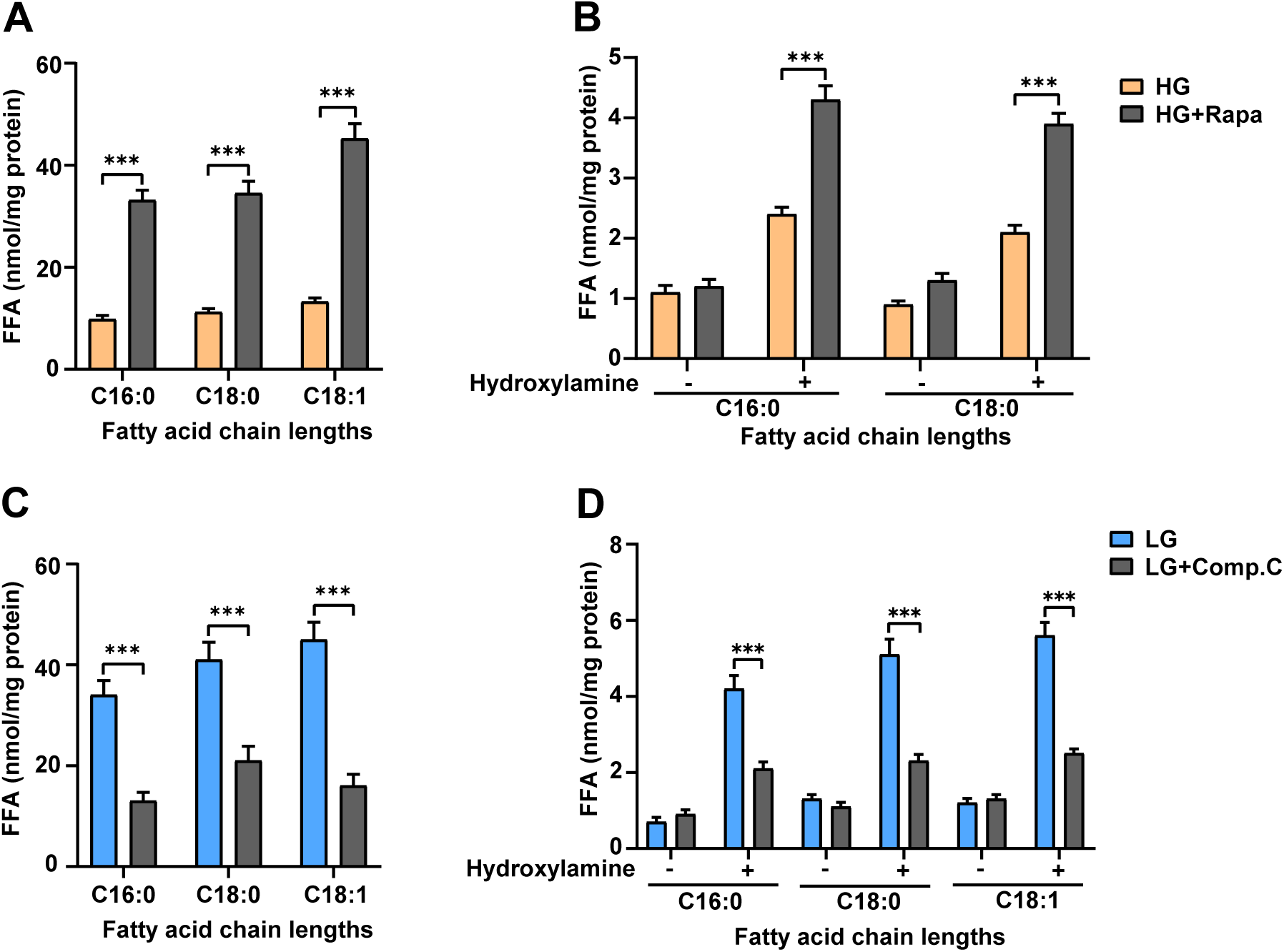
Anabolic and catabolic factors impinge on fatty acylation of proteins. **(A and B)** Primary hepatocytes grown in the presence of mTOR inhibitor, rapamycin (N=2, n=3). Fatty acids assayed from (A) cellular pools and (B) membrane fractions subjected to hydroxylamine treatment. **(C and D)** Primary hepatocytes treated with Compound C (Dorsomorphine), an AMPK inhibitor, and fatty acids analyzed (N=2, n=3) from (C) cellular pools and (D) their respective membrane fractions. Data are represented as mean ± SEM and analyzed by the Student’s t-test and ANOVA. A value of *p* ≤ 0.05 was considered statistically significant. \*\*\**p* ≤ 0.001.

The intracellular FFA concentration in a fasting state is also dependent upon mitochondrial β-oxidation. Therefore, we asked if directly inhibiting CPT1, which is essential for β-oxidation, affected fatty acylation under fasted conditions. Interestingly, incubating cells with etomoxir, a known inhibitor of CPT1, led to a significant decrease in protein fatty acylation (**Figure 7A**). Even though it was surprising to see this dependence on mitochondrial fatty acid utilization, this was well corroborated by the reduction in intracellular FFA pools (**Figure 7B and Supplementary Figure S6C**). Collectively these results indicated the central role of factors that directly affect fatty acid uptake and utilization, and mitochondrial functions in mediating fatty acylation (**Figure 7C)**.

**Figure 7:**
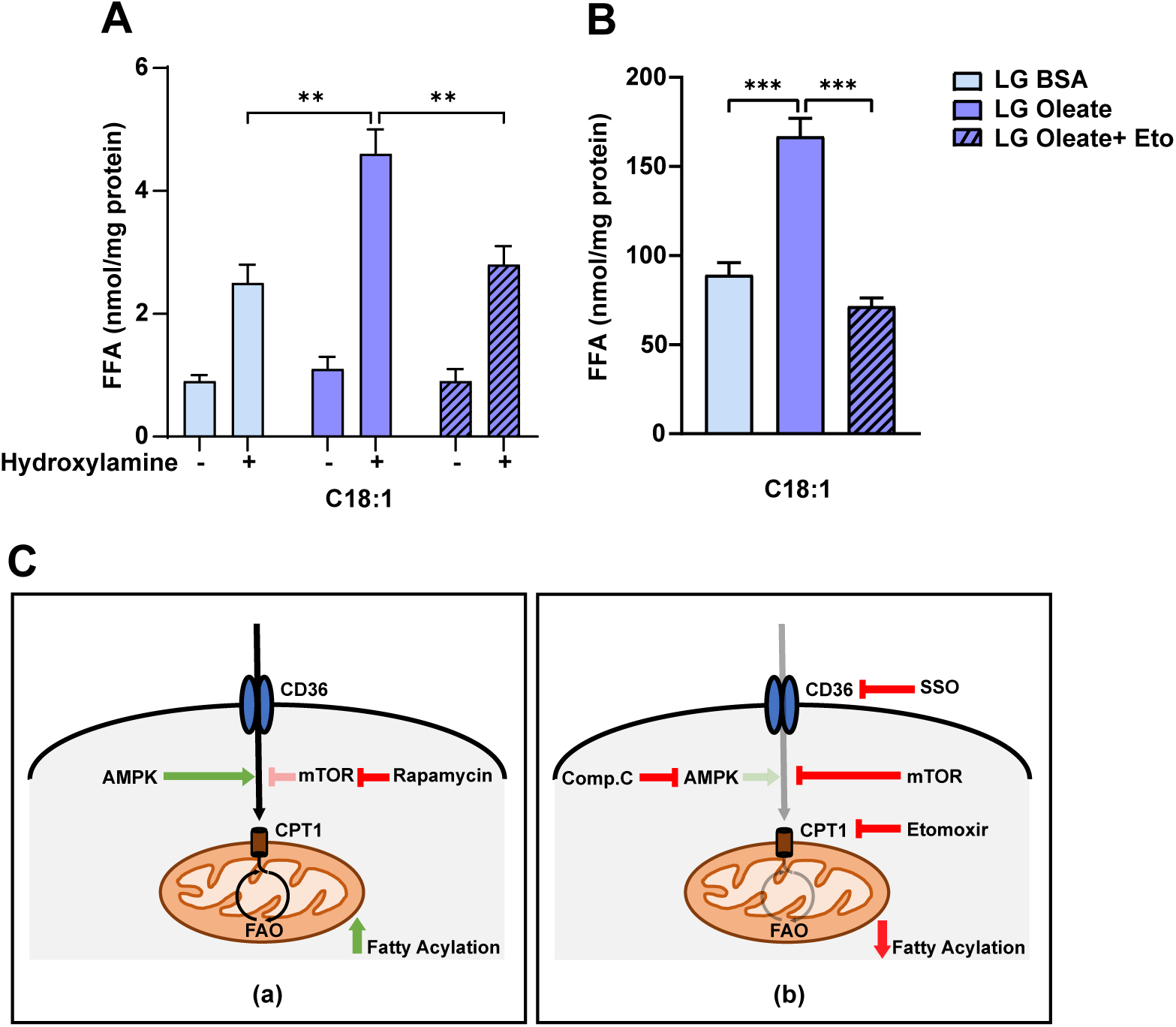
Impact of mitochondrial fatty acid transport on fatty acylation. Primary hepatocytes grown in the presence of BSA or BSA conjugated oleate in the presence of CPT-1 inhibitor, etomoxir (N=2, n=3). **(A)** Fatty acid release assessed from membrane fractions post hydroxylamine treatment and **(B)** Corresponding cellular FFA pools. **(C)** Schematic representation of the interplay of anabolic, catabolic, and mitochondrial regulations modulated using chemical inhibitors and genetic perturbations impinging on regulatory kinases and thereby the protein fatty acylation. Data are represented as mean ± SEM and analyzed by the Student’s t-test and ANOVA. A value of *p* ≤ 0.05 was considered statistically significant. \*\*\**p* ≤ 0.001.

### SIRT4-dependent mitochondrial energetics and metabolism exert control over fatty acylation

Our studies thus far hinted that fed-fast-dependent change in global protein acylation also involved aspects of mitochondrial metabolism. This prompted us to investigate whether any mitochondrial factors could drive this coupling. We and others have demonstrated that SIRT4, a mitochondrial NAD-dependent deacylase is an essential component that regulates β-oxidation and cellular energetics in a fed-fast dependent manner (34, 46–49). Thus, we next addressed the role of SIRT4, which would also provide additional evidence for mitochondrial dependence for the phenomena. C16-alkyne-based click-chemistry in cells stably overexpressing SIRT4 showed a significant decrease in protein palmitoylation in both fluorescence imaging (**Figure 8A**) and gel-based fluorescence assays **(Figure 8B)**. Interestingly, while exogenous expression of SIRT4 decreased the extent of fatty acylation in cells grown under LG conditions, liver lysate of mice devoid of SIRT4 (SIRT4-KO) displayed the opposite effect (**Supplementary Figure S7A**). We next wanted to assess the quantitative changes in the FFA pool and associated protein fatty acylation in genetic perturbations of SIRT4. Towards this, we used primary hepatocytes isolated from SIRT4KO mice, and then using the adenoviral technology, transduced either a control (Ad-CMV) vector or SIRT4 rescue (Ad-SIRT4) constructs (**Supplementary Figure S7B**). As anticipated, we found a substantial decrease in intracellular FFA pools, independent of the chain length or unsaturation, in hepatocytes devoid of SIRT4 after rescue with adenoviral SIRT4 (**Supplementary Figure S7C**). This also revalidated the central role that SIRT4 plays in regulating lipid metabolism in hepatocytes. More importantly, on measuring both soluble and protein-released FFAs from the same samples, we found a consistent reduction in hydroxylamine-sensitive fatty acylation when SIRT4 was rescued in SIRT4 devoid hepatocytes (**Figure 8C**, **8D**). Further, we also observed SIRT4-dependent dampening in FFA levels in cells that were fed with increasing amounts of fatty acid, as indicated in **Supplementary Figure S7D.**

**Figure 8:**
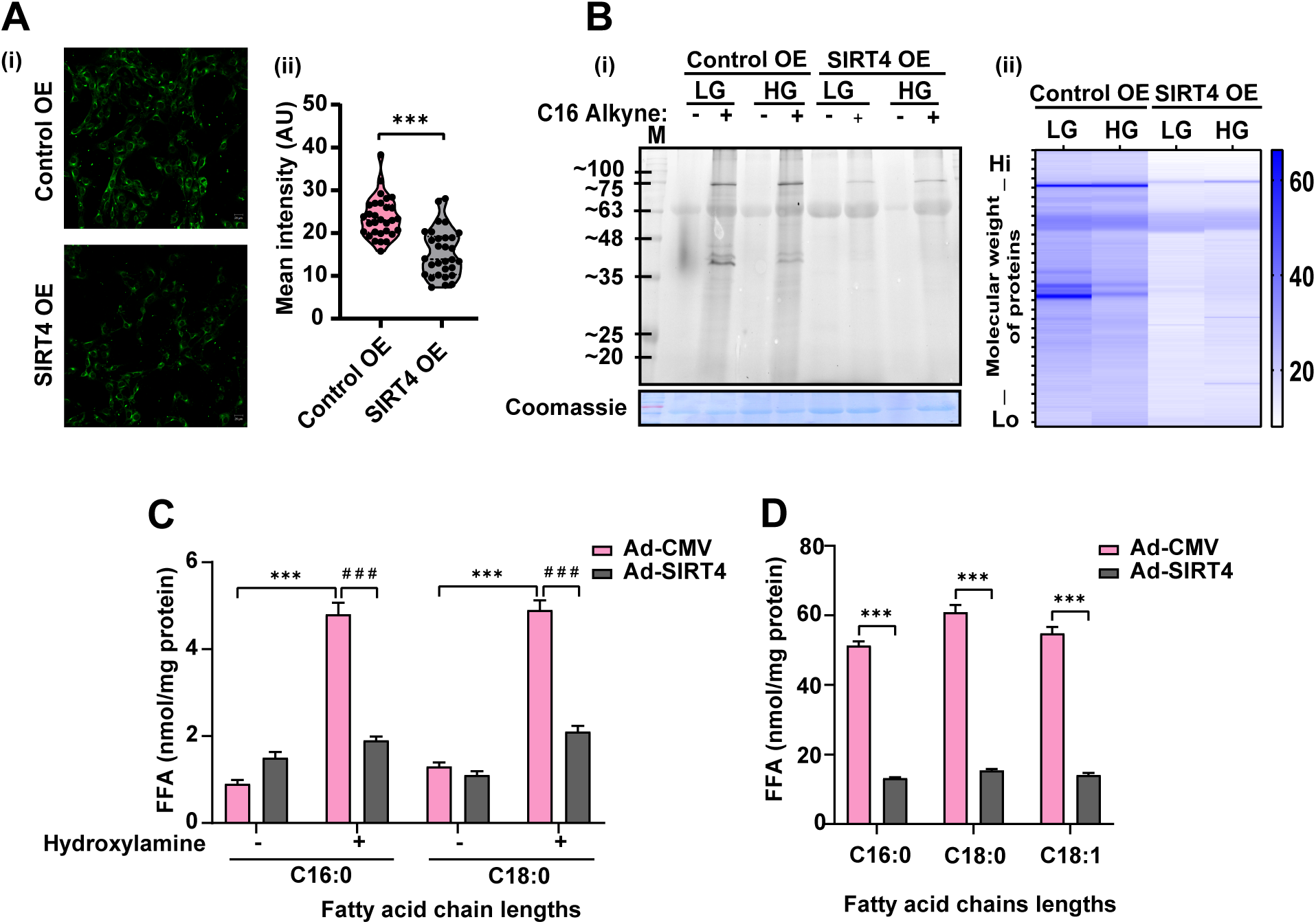
Mitochondrial sirtuin SIRT4 regulates global S-fatty acylation. **(A)** NIH 3T3 cells stably overexpressing control and SIRT4 were grown under LG and HG conditions in the presence of C16-alkynyl FA for 4 hours. Cells were fixed, permeabilized, and reacted with Alexa Fluor 488 azide (N=2, n=30). (i) Imaging was done using confocal microscopy (scale bar 20 microns), (ii) Quantification of fluorescence intensity. **(B)** Click chemistry-based analysis of protein palmitoylation in control and SIRT4 overexpressing HEK293T stable cells subjected to C16-alkyne treatment in LG and HG-containing media, DMSO was used as control (N=2), (i) Visualized by in-gel fluorescence, (ii) Heat map depicting band intensities in respective lanes. **(C)** Free fatty acids estimated post-release using hydroxylamine treatment in SIRT4 devoid hepatocytes with and without Ad-SIRT4 overexpression (N=3, n=3). **(D)** Estimate of cellular FFA pools extracted from SIRT4 devoid hepatocytes post overexpressing hSIRT4 (N=3, n=3). Data are represented as mean ± SEM and analyzed by the Student’s t-test and ANOVA. A value of *p* ≤ 0.05 was considered statistically significant. \*\*\**p* ≤ 0.001; ^###^*p* ≤ 0.001.

Given the multiple modes of SIRT4-dependent control of fatty acid utilization and metabolism, we set out to delineate if the effects we observed were contributed by cellular uptake and/or β-oxidation. Towards this, the aforementioned SIRT4 devoid hepatocytes rescued by adenoviral SIRT4 were treated with SSO or etomoxir to inhibit CD36-mediated fatty acid uptake and CPT1-mediated β-oxidation, respectively. While SSO treatment led to a significant decrease in soluble FFA pools in SIRT4-KO hepatocytes (**Figure 9A**, Supplementary Figures S8A **and S8B**), which was expected, interestingly, inhibition of CPT1 also led to a similar response (Supplementary Figure S8C). Although counterintuitive and intriguing, this pointed towards the mitochondrial β-oxidation mediated regulation of fatty acid uptake whose mechanistic details need further investigation. Nonetheless, since CD36-dependent FFA uptake emerged as consistently being rate limiting vis-à-vis protein fatty acylation, we asked if enhanced modification in the absence of SIRT4 was sensitive to SSO treatment. Here, we found that inhibiting CD36 decreased hydroxylamine sensitive protein conjugated fatty acids (irrespective of the chain length and unsaturation (**Figure 9B**, Supplementary Figures S8D **and S8E**)) in SIRT4 null hepatocytes but not in those wherein SIRT4 expression was rescued.

**Figure 9:**
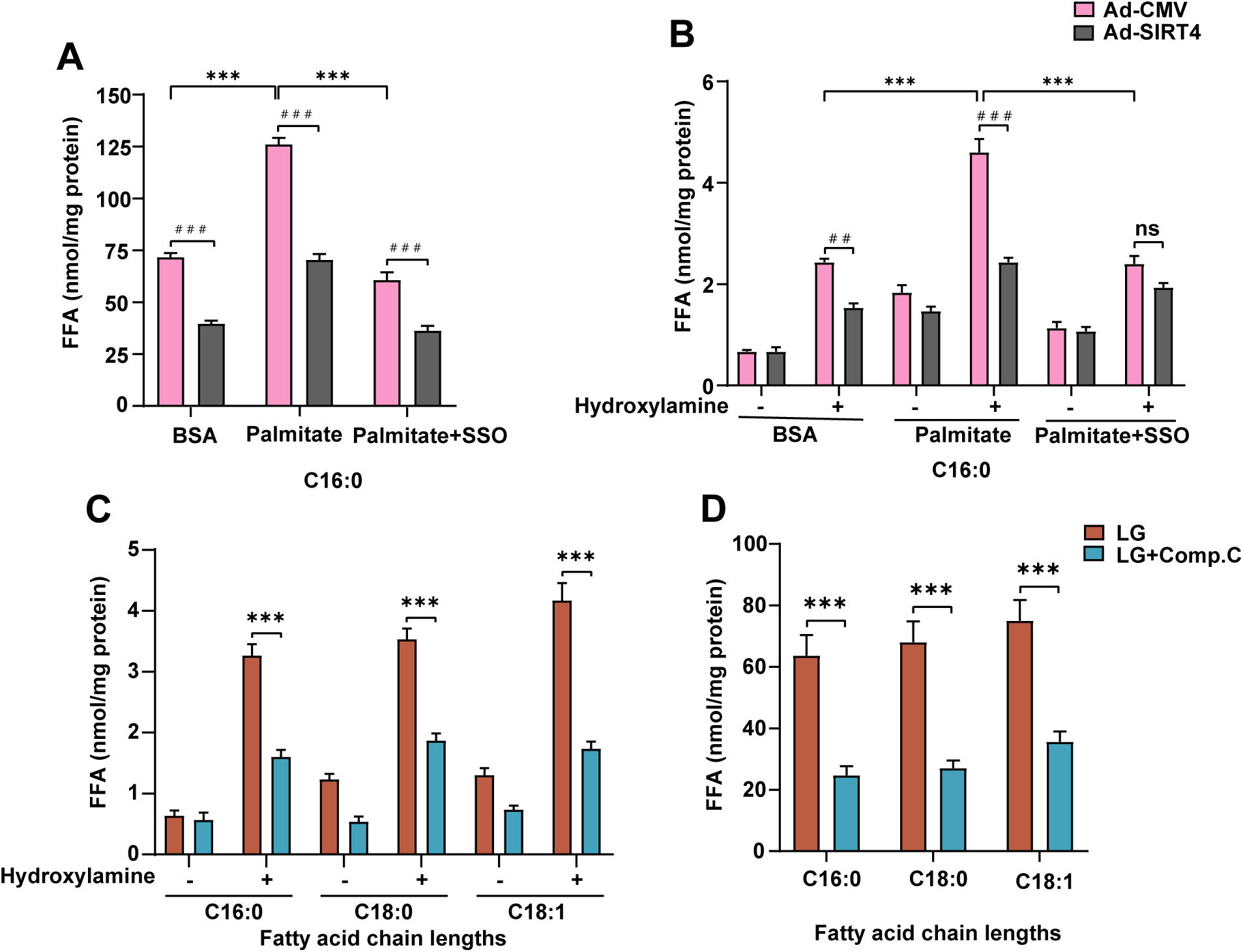
Absence of SIRT4 mimics fasted state and leads to increased protein acylation. **(A and B)** SIRT4 null hepatocytes transduced with either Ad-CMV or Ad-SIRT4 grown in LG media were subjected to treatment with SSO, CD36 inhibitor (N=2, n=3). (A) Their cellular pools were assayed for free fatty acid content and (B) Membrane fractions were analyzed for fatty acid release post-hydroxylamine treatment. **(C and D)** Primary hepatocytes from SIRT4 null mice were treated with Compound C, an AMPK inhibitor in low glucose (N=2, n=3), (C) fatty acid release analyzed from membrane fractions post hydroxylamine treatment and (D) respective cellular fraction. Data are represented as mean ± SEM and analyzed by the Student’s t-test and ANOVA. A value of *p* ≤ 0.05 was considered statistically significant. \*\*\**p* ≤ 0.001; ^##^*p* ≤ 0.01; ^###^*p* ≤ 0.001.

The ability of SIRT4 to exert control over lipid metabolism also involves antagonistic interplay with AMPK that is brought about by changes in mitochondrial energetics (46). Thus, we addressed the role of the SIRT4-AMPK axis, which was also motivated by our results that implicated AMPK as a key determinant of global fatty acylation. Consistent with our hypothesis, increased fatty acylation seen in SIRT4 null hepatocytes was significantly lowered following AMPK inhibition (**Figure 9C**), and this correlated well with decrease observed in soluble FFA pools (**Figure 9D**). Together these findings suggested the involvement of mitochondrial energetics and SIRT4-AMPK dependent retrograde signaling in governing cellular FFA homeostasis and consequential covalent modification of proteins.

## DISCUSSION

It is intuitive to expect post-translational modifications of proteins to have bidirectional coupling with cellular physiology. This is especially relevant in contexts wherein concentrations of intracellular metabolite, dictated by both its uptake and utilization for anabolic/catabolic processes and energetics, determine their availability for such post-translational modifications. In this regard, while protein fatty acylation has been established amongst key regulatory post-translational modifications, whether it is governed by cellular metabolic and energetic status is not well understood. In this context, here, we provide substantial evidence to highlight the pivotal role of glucose-dependent metabolic shifts and mitochondrial functions towards impacting global protein fatty acylation. In addition to this, in conditions that mimic fed and fasted states, we have also elucidated this dependence on metabolic sensing.

Fatty acids are covalently linked to proteins via amide linkages at lysine (*N*-linked) or thioester bonds at cysteine (*S*-linked) residues (27). Even though fatty acid chain length-dependent bias is not well established, protein fatty acylation mediated by > C16 has been proposed to be largely through thioester binds with protein cysteine residues (20, 21). In this context, our study has utilized complementary approaches, such as alkyne-based click chemistry, acyl biotin exchange, and quantitative LC-MS/MS-based lipidomics to score for the global changes in protein fatty acylation across physiological states. To assess fatty acylation by both the gel and lipidomics-based assays, we have employed hydroxylamine treatment, a method that is well established and has been shown to preferentially target *S*-linked modifications (41). Our results, across physiological and pharmacogenetic perturbations, clearly illustrate high concordance between these approaches. Moreover, hydroxylamine-sensitive global fatty acylation seems to be mostly contributed by long-chain fatty acids.

Fatty acylation of proteins has been shown to regulate several physiological processes, even in evolutionarily diverse organisms. Besides altering localization by providing membrane anchorage, fatty acylation of membrane-bound/embedded proteins is also known to regulate their structure and function (8, 9). Despite these findings, whether this is dictated by intracellular concentrations of FFAs has not been well documented. In this regard, using exogenously added fatty acids, we have found a concentration or uptake dependence that drives the extent of fatty acylation. More importantly, by addressing this in primary hepatocytes especially under conditions that show dynamic changes in intracellular FFA levels, we provide a physiological setting for substrate-dependent protein modification, as discussed below.

Intracellular pools of FFAs in hepatocytes are determined by flux through lipogenesis, lipolysis, and β-oxidation. It is important to note that these processes are in turn regulated by anabolic and catabolic demands during fed and fasted conditions. Specifically, the availability of glucose under these states is known to dictate the FFA pool inside cells. Therefore, even though glucose-dependent fatty acid metabolism has been well studied, whether that impinges on global protein fatty acylation has not been addressed thus far. In this regard, here, we clearly illustrate that cells grown in media containing HG or LG show robust changes in hydroxylamine-sensitive protein fatty acylation. To reiterate, these findings indicate a cross-talk between two key nutrient sources, and as to how the availability of carbohydrates influences post-translational modification brought about by fatty acids. Moreover, this will likely have an impact on evaluating the impact of glucose metabolism on protein fatty acylation under various physiological and pathological states.

To provide a mechanistic basis for fatty acid flux dependence, we perturbed cellular uptake and mitochondrial β-oxidation by inhibiting CD36 and CPT1, respectively. While the role of CD36 was anticipated, we have found that inhibiting mitochondrial uptake/utilization via CPT1 has a comparable effect vis-à-vis both soluble FFA pools and hydroxylamine-sensitive protein fatty acylation status. Mitochondrial fatty acid oxidation is coupled to metabolic sensing via AMPK, Sirtuins, and mTOR, which collectively also govern cellular physiology in fed and fasted states. In this regard, our results demonstrate the involvement of AMPK and TOR signaling and also provide new evidence on potential unexplored mechanisms that operate in fed and fasted states in determining the extent of fatty acylation. We and others have previously shown the role of SIRT4 in cellular physiology and mitochondrial fatty acid metabolism. Consistent with this, SIRT4 which mediates an anabolic state, downregulates global fatty acylation. Given that we have recently found the pivotal role of SIRT4 in coupling mitochondrial metabolism/energetics and AMPK/TOR signaling, which orchestrate fed and fasted physiology (47), we investigated if this axis impinges on protein fatty acylation. It is important to note that our findings, using loss-/gain-of-function perturbations and genetic rescue of SIRT4, demonstrate its role in dictating hydroxylamine-sensitive conjugation of fatty acids to proteins. Further, SIRT4 perturbations mimic global protein fatty acylation changes that were observed in fed and fasted conditions. As hypothesized, based on previously described functions of SIRT4, we also found that this is intricately linked with mitochondrial energetics. Notably, combinatorial chemogenetic perturbations employing wild type and SIRT4-null hepatocytes and treatments with activators/inhibitors that affect β-oxidation, including AMPK and CPT1, have provided confirmatory evidence. Therefore, besides validating the interplay with cellular physiology, our study here provides further mechanistic insights into nutrient inputs-dependent modification of proteins that is mediated by fatty acids.

## EXPERIMENTAL PROCEDURES

### Animals used

All animal experiments were performed in accordance with the Institutional Animal Ethics Committee (IAEC) guidelines. C57BL/6 and SIRT4 knockout mice (obtained from Jackson Laboratories, catalog no. 012756) were used for experiments. In the fed-starved experiments, *ad libitum*-fed mice were used as the fed cohort, and for the fasted group mice were starved for 24 hours. Liver tissues were collected and snap-frozen in liquid nitrogen.

### Cell lines

HEK293, HEK293T, and C2C12 cells were cultured as recommended by ATCC. Cells were grown either in high glucose (25 mM) (HG) (Sigma-Aldrich, catalog no. D7777) or low glucose (5 mM) (LG) (Sigma-Aldrich, catalog no. D5523) containing DMEM media as indicated.

### Primary hepatocyte culture

Hepatocytes were isolated from C57BL/6 and SIRT4 KO mice (8–12 weeks old) by perfusing the liver tissue with collagenase type IV (Sigma-Aldrich, catalog no. C5138) containing media as described in Shaw et al. (47). Cell pellet was suspended in DMEM HG media having 10% (v/v) FBS (Gibco, catalog no. 16000-044), counted and plated in collagen-coated plates (5 µg/cm^2^) (Sigma-Aldrich, catalog no. C3867). Cells were allowed to adhere for 6 hours and media was changed to incomplete DMEM HG. Cells were collected post 48–72 hours of plating after respective treatments.

### Plasmids, transfections, and transduction

pBabe puro with/without hSIRT4-HA-Flag was used for transfection to stably overexpress SIRT4 in cell lines using Lipofectamine 2000 (Invitrogen, catalog no. 11668019) following the manufacturer’s protocol. SIRT4 overexpression in primary hepatocytes was done using an adenoviral overexpression system as described in Shaw et al. (47). The supernatant (Adenovirus) was added 16–18 hours post cell plating for viral transduction. Media was changed after 18–20 hours of virus addition. Metabolic shifts and pharmacological treatments were done as described below.

### Drug treatments

HG and LG treatments were given for 12 hours in media containing 10% (v/v) FBS. For HG, media was replenished 3 hours before cell collection. For mTOR inhibition and AMPK activation, 20 nM Rapamycin (Sigma-Aldrich, catalog no. R0395) and 0.5 mM AICAR (Sigma-Aldrich, catalog no. A9978) treatments were given in complete HG media for 6 hours, respectively. 20 µM Compound C (Dorsomorphine, Sigma, catalog no. A5499) treatment was given in LG media for 6 hours for AMPK inhibition. To induce uncoupling in cells, 0.5 µM FCCP (Sigma-Aldrich, catalog no. C2920) treatment was given in complete HG media for 12 hours. To inhibit the activity of PATs, 150 µM 2-bromopalmitate treatment (Sigma-Aldrich, catalog no. 21604) was given for 16 hours in complete LG media.

### Fatty acid uptake assay

Here, 100 µM, 250 µM, and 500 µM palmitate (Sigma-Aldrich, catalog no. P9767) and oleate (Fisher Scientific, catalog no. 75096) conjugated to fatty acid-free BSA were used for feeding cells in incomplete LG media. Fat-free BSA was used as a control. Etomoxir treatment (Sigma-Aldrich, catalog no. E1905) was given at a concentration of 60 µM for 16 hours to inhibit CPT1. Sulfo-N-succinimidyl oleate sodium (SSO) (Sigma-Aldrich, catalog no. SML2148) was used to inhibit CD36 at a concentration of 50 µM for 12 hours. Fatty acid uptake was detected using LC-MS/MS as subsequently described.

### Acyl exchange assay

Total protein fatty acylation was detected using a modification in the acyl biotin exchange protocol (50). Protein lysates were prepared in RIPA buffer (50 mM Tris HCl pH 7, 150 mM NaCl, 1% Triton X-100, 0.5% w/v Sodium deoxycholate, 0.1% SDS). 80– 100 µg of lysate was taken per sample and subjected to 50 mM *N*-ethylmaleimide (Catalog no. E3876) treatment for 2 hours on ice to block the free cysteines in proteins. The sample was distributed equally into 2 tubes and proteins were precipitated out overnight using chilled acetone at –20 °C. One sample was treated with 2 M hydroxylamine, pH 7.2 (NH_2_OH, Hydroxylamine hydrochloride, Sigma-Aldrich, catalog no: 159417) for 2 hours on a shaker at 25 °C. The control sample was left untreated (-HAM). Hydroxylamine leads to the selective removal of *S*-linked acyl chains from proteins. Proteins were precipitated overnight and the samples, both ± HAM, were re-suspended in lysis buffer to be treated with 0.5 µM rhodamine red C2 maleimide for 2 hours on ice (Invitrogen, catalog no. R6029). The proteins were then precipitated overnight with chilled acetone, pelleted down and visualized using SDS-PAGE.

### Metabolic labeling with C16 alkyne

Alkynyl palmitic acid used for metabolic labelling was procured from Sigma Aldrich (catalog no. 900400P) and the protocol was followed as described (51). To assay palmitoylation, 100 µM C16-alkyne was conjugated with fatty acid-free BSA (5% v/v) (Sigma-Aldrich, catalog no. A7030) in DMEM LG or HG media by sonication (15 min) followed by incubation at room temperature (15 min). Cells were treated in respective media for 18–24 hours. Only DMSO (Sigma-Aldrich, catalog no. D2650) was added as vehicle control.

### Click chemistry

Protein lysates from C16-treated cells were prepared using a lysis buffer and detected using Click chemistry (51). C16-alkynes bound to proteins were conjugated to the fluorescent azide in a copper-catalyzed Huisgen cycloaddition reaction. 30–50 µg of protein lysate was treated with AlexaFluor 488 azide (Invitrogen, catalog no. A10266) in the presence of TCEP (Sigma-Aldrich, catalog no. C4706), TBTA (Sigma-Aldrich, catalog no. 678937), CuSO_4_ and in the dark, at room temperature for 1 hour. The reaction was terminated by precipitating the protein overnight, in chilled acetone. Samples were assessed using SDS-PAGE.

### Detection/visualization of *S*-acylation on gel

Samples boiled in Laemmli buffer were run on polyacrylamide gels containing 12–15% resolving and 4% stacking. Gels were imaged using Typhoon 7000 (GE Healthcare). C16-alkyne samples labelled with AlexaFluor 488 azide using click chemistry and Rhodamine red C2 maleimide labelled using acyl exchange were visualized at 488 nm and 548 nm, respectively. Imaging at 647 nm was done to visualize the protein ladder. Gels were then either stained using Coomassie staining solution for protein normalization or samples were transferred onto PVDF membrane by western blotting, blocked in 5% skimmed milk, and probed with antibody for β-actin (catalog no. A1978) or β-tubulin (catalog no. T8328) as loading controls, both from Sigma Aldrich.

### Immunofluorescence

C16 alkyne treatment was given to cells grown on coverslips, as stated previously, for 4 hours. Media was removed from wells and coverslips were washed in PBS. Fixation was done using 4% PFA for 10 min. Cells were then permeabilized using 0.1% Triton X100 for 5 min at room temperature. Click reaction was done on the coverslips using AlexaFluor 488 azide in a 100 µL reaction volume containing TCEP and CuSO_4_ in PBS (52). The reaction was carried out in the dark at room temperature. 6–8 washes of PBS were given for each coverslip to remove excess stain. Coverslips were further mounted using an anti-fade solution. Imaging was done using a Zeiss 510 confocal microscope.

### Fatty acid release assay and LC-MS

Cells were collected post-respective treatments, washed in PBS, pelleted down, and stored at –80 °C until further processing. Pellets were resuspended in PBS by probe sonication (water sonication used in adenovirus-containing samples). Suspensions were then centrifuged at 100,000 g for 40 min at 4 °C. The supernatant was used as the cell soluble fraction. The pellet (membrane fraction) was further used for delipidation by treating it with the methanol-chloroform solution as reported earlier (53, 54). The protein layer was collected and washed in methanol:chloroform: PBS solution and was resuspended in PBS using probe sonication. The amount of protein was quantified using a BCA protein estimation kit (Pierce). Samples were divided into two for treatment with and without hydroxylamine for 2 hours at 25 °C on a shaker. The reaction was stopped using methanol:chloroform in a 1:2 ratio spiked with C17:1 FFA (1 nmol as an internal standard). The organic layer was collected and dried in a nitrogen stream. FFAs were detected using LC-MS/MS using protocols previously reported by us (55–57). Values were normalized to internal control 17:1 FFA and the respective protein concentrations.

### RNA isolation and PCR

Total cellular RNA was extracted using Trizol (catalog no. 15596018) as per the manufacturer’s instructions. 1 µg of total RNA was used for cDNA preparation using SuperScript IV reverse transcriptase kit (Catalog no. 18090200). PCR was done using KAPA SYBR® FAST Universal 2X qPCR Master Mix (Catalog no.). The following primer pairs were used for assaying overexpression of human SIRT4, forward (5′-CAGCGCTTCATCACCCTTTC-3′) and reverse (5′-CCTACGAAGTTTCTCGCCCA-3′); and normalized to mouse actin B, forward (5′-GCATGGGTCAGAAGGATTCC-3′) and reverse (5′-ACGCAGCTCATTGTAGAAGG-3′).

### Data processing and analysis

In-gel fluorescence, western blot, and cell immunofluorescence images were analyzed using the ImageJ software. Student’s t-test and analysis of variance (ANOVA) were used for statistical analysis. Data were processed in Microsoft Excel, and the statistical significance was calculated using Excel or GraphPad Prism. The results are expressed as means ± standard deviations (SD)/standard error (SE) as indicated. All experiments were performed at least twice (N) with a minimum of three to four replicates (n).

### Supporting information

This article contains supporting information

## Acknowledgments

We thank Dr. Kalidas Kohale and Dr. Shital Suryavanshi (TIFR-AH) and Dr. Suraj Ingle, Dr. Sagar Tarte, Ms. Ritika Gupta, Mr Chetan (National Facility for Gene Function in Health and Disease, IISER Pune) for help with the animal experiments. We thank Hema P. Bagul for her help with the C16-alkyne experiment. We thank Dr. Abinaya Rajendran, Alaumy Joshi and Saddam Shekh for technical assistance with the LC-MS/MS experiments.

## Funding and additional information

This study was supported by funds from DAE/TIFR-Govt. of India (Grant number P0116) to U.K., Swarnajayanti Fellowship (DST Govt. of India. Grant number DST/SJF/LSA-02/2012-13) to U.K., SwarnaJayanti Fellowship from the Science and Engineering Research Board (SERB), Department of Science and Technology (DST), Government of India (Grant number SB/SJF/2021-22/01) to S.S.K., and the Department of Science & Technology–Funds for Improvement of S&T Infrastructure Development (DST-FIST) (grant number SR/FST/LSII-043/2016) to the Department of Biology, IISER Pune. We also thank the National Facility for Gene Function in Health and Disease (supported by a grant from the Department of Biotechnology, Government of India; BT/INF/22/ SP17358/2016) at IISER Pune for maintaining and providing mice used in this study. A.C. thanks a Graduate Student Fellowship from the Council for Scientific and Industrial Research (CSIR).

---

**Supplementary Figure 1:** Acyl exchange done using protein lysates where free cysteines are blocked using N-ethyl maleimide (NEM), subjected further to fatty acid release using KOH or hydroxylamine (HAM) and detected using Rhodamine C2 maleimide to detect free cysteines post HAM treatment. Rhodamine C2 maleimide was done in the untreated sample as seen in lane 2. Corresponding Coomassie-stained gel for loading control.

**Supplementary Figure 2:** Total free fatty acid (FFA) composition of primary hepatocytes grown in high glucose (HG) and low glucose (LG) media (n=3). Data are represented as mean ± SEM and analyzed by the Student’s t-test. A value of *p* ≤ 0.05 was considered statistically significant. \*\*\**p* ≤ 0.001.

**Supplementary Figure 3: (A)** Primary hepatocytes were grown in the presence of either BSA or in increasing concentration of BSA conjugated palmitate (n=3). **(B and C)** Primary hepatocytes grown in the presence of BSA and BSA conjugated palmitate subjected to treatment with SSO (N=2, n=3), (B) showing the release of stearate (C18:0) and (C) oleate (C18:1), post hydroxylamine treatment. **(D)** Showing corresponding levels of stearate (C18:0) and oleate (C18:1) in cellular pools. Data are represented as mean ± SEM and analyzed by the Student’s t-test and ANOVA. A value of p ≤ 0.05 was considered statistically significant. \*\**p* ≤ 0.01; \*\*\**p* ≤ 0.001.

**Supplementary Figure 4:** Estimate of C18:0 and C18:1 from primary hepatocytes grown in LG medium and treated with 2-BP (N=2, n=3), **(A)** Membrane fraction after hydroxylamine induced fatty acid release and **(B)** the respective cellular pools. Data are represented as mean ± SEM and analyzed by the Student’s t-test and ANOVA. A value of *p* ≥ 0.05 was considered statistically non-significant (ns).

**Supplementary Figure 5: (A) (i)** Representative western blots showing pS6K^T389^, total S6K, and actin for primary hepatocytes treated with rapamycin **(ii)** Quantitation of pS6K^T389^/S6K normalized to respective actin blots (N=2, n=3). **(B) (i)** Representative western blots showing pAMPK^T172^, total AMPK, and actin for primary hepatocytes treated with Compound C (Dorsomorphine) **(ii)** Quantitation of pAMPK^T172^/AMPK normalized to respective actin blots (N=2, n=3). Data are represented as mean ± SEM and analyzed by the Student’s t-test. A value of *p* ≤ 0.05 was considered statistically significant. \*\**p* ≤ 0.01.

**Supplementary Figure 6: (A) (i)** Representative western blots showing pAMPK^T172^, total AMPK, and actin for primary hepatocytes treated with AICAR **(ii)** Acyl exchange for protein acylation using protein lysates from primary hepatocytes grown in DMEM media containing HG with or without AICAR treatment **(iii)** Heat map depicting mean of band intensities in respective lanes (N=2, n=3). **(B) (i)** Representative western blots showing pAMPK^T172^, total AMPK, and actin for primary hepatocytes treated with FCCP, **(ii)** Primary hepatocytes were treated with FCCP in HG conditions, proteins were subjected to acyl exchange, with and without hydroxylamine and visualized on a gel, **(iii)** Heat map depicting mean of band intensities in respective lanes (n=3). **(C)** Fatty acid content in primary hepatocytes post-feeding of increasing concentrations of BSA conjugated oleate in the presence and absence of etomoxir, only BSA was used as control (n=3). Data are represented as mean ± SEM and analyzed by the ANOVA. A value of *p* ≤ 0.05 was considered statistically significant. \**p* ≤ 0.05; \*\**p* ≤ 0.01; \*\*\**p* ≤ 0.001; ^#^*p* ≤ 0.05 (* represents statistical differences between group LG BSA and LG Oleate; ^#^ represents statistical differences between group LG Oleate and LG Oleate + Eto)

**Supplementary Figure 7: (A) (i)** Total protein fatty acylation observed using acyl exchange in liver tissue isolated from isogenized wild type and SIRT4 null mice, lysates were treated with NEM and HAM and further detected in gel using Rhodamine maleimide. The coomassie-stained gel was used for depicting loading control (n=3) **(ii)** Heat map depicting mean of band intensities in respective lanes. **(B)** Representative image of PCR showing hSirt4 amplification post adenoviral overexpression of SIRT4 in SIRT4 devoid hepatocytes. **(C)** Total free fatty acid content of primary hepatocytes from SIRT4 null mice grown in LG media post-restoration of SIRT4 expression using adenoviral transduction (n=3). **(D)** Palmitate uptake assayed in SIRT4 null hepatocytes post Ad-SIRT4 overexpressed with increasing concentration of BSA conjugated palmitate (n=3). Data are represented as mean ± SEM and analyzed by the Student’s t-test and ANOVA. A value of *p* ≤ 0.05 was considered statistically significant, \*\*\**p* ≤ 0.001.

**Supplementary Figure 8: (A-D)** Primary hepatocytes from SIRT4 KO mice grown in the presence of BSA and BSA conjugated palmitate subjected to treatment with SSO (N=2, n=3), Estimate of **(A)** stearate and **(B)** oleate in soluble pools and **(D)** stearate and **(E)** oleate, post hydroxylamine treatment in the respective membrane pool. **(C)** Fatty acid uptake assayed in SIRT4 null hepatocytes with increasing concentration of BSA conjugated oleate in LG conditions, inhibited by etomoxir, a CPT-1 inhibitor (n=3). Data are represented as mean ± SEM and analyzed by the Student’s t-test and ANOVA. A value of *p* ≤ 0.05 was considered statistically significant. \**p* ≤ 0.05; \*\**p* ≤ 0.01; \*\*\**p* ≤ 0.001; ^#^*p* ≤ 0.05; ^##^*p* ≤ 0.01; ^###^*p* ≤ 0.001.

