## Supplementary figures and images for "Metabolic transitions regulate global protein fatty acylation"

### Supplementary information and figures

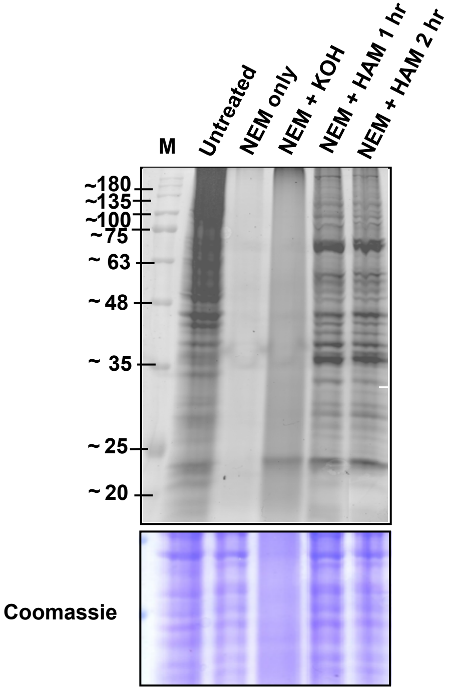

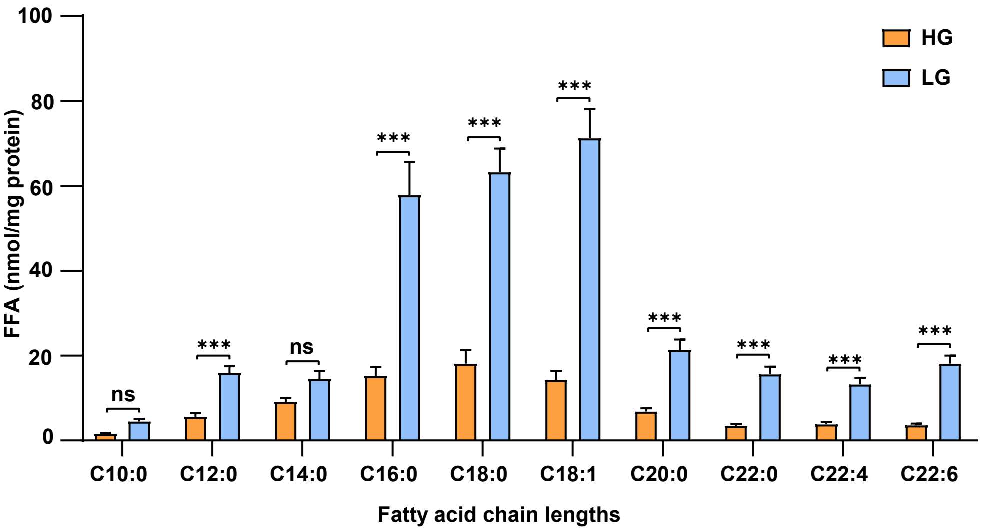

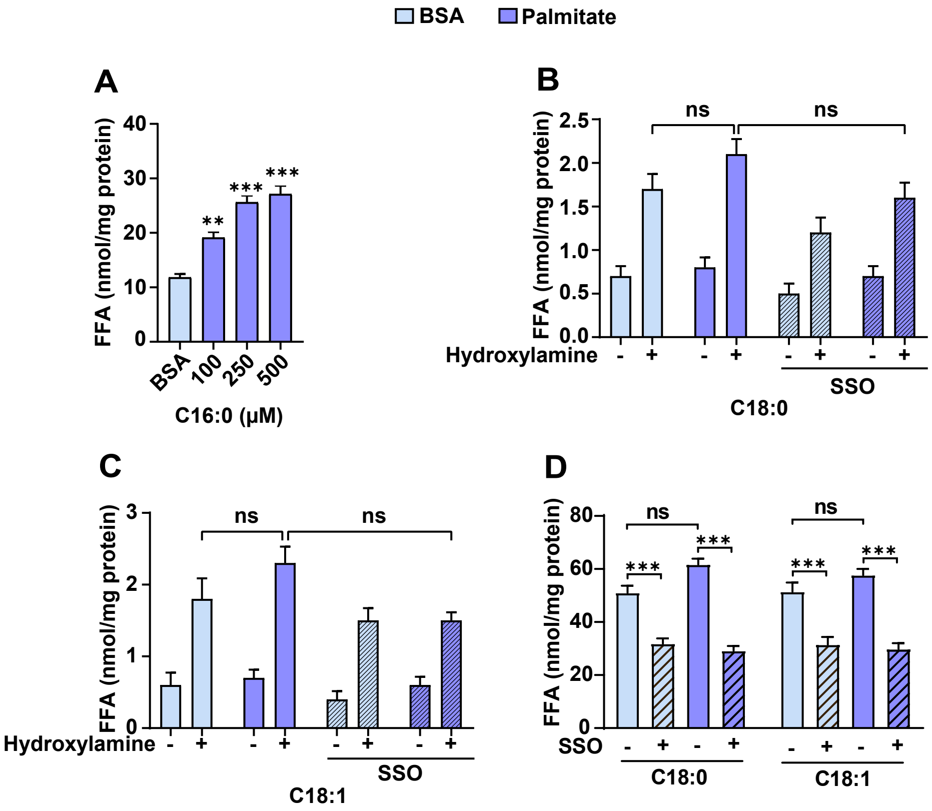

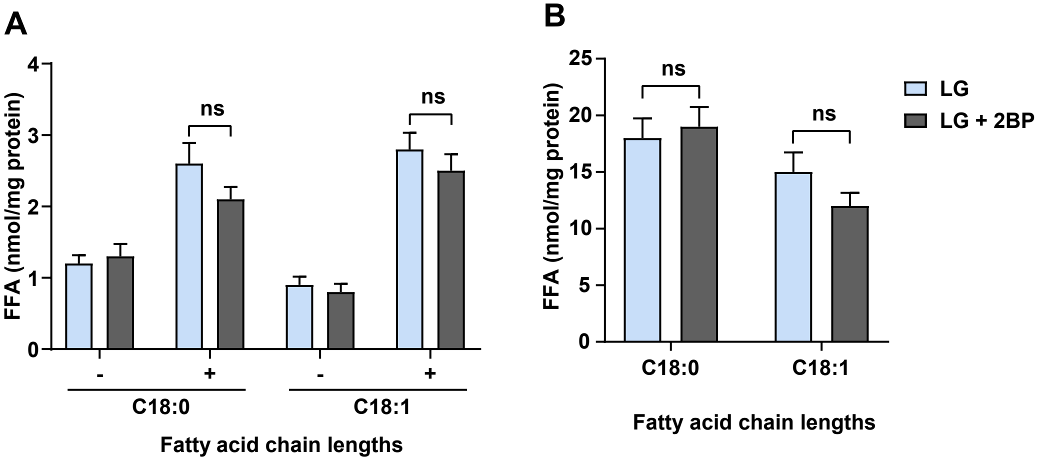

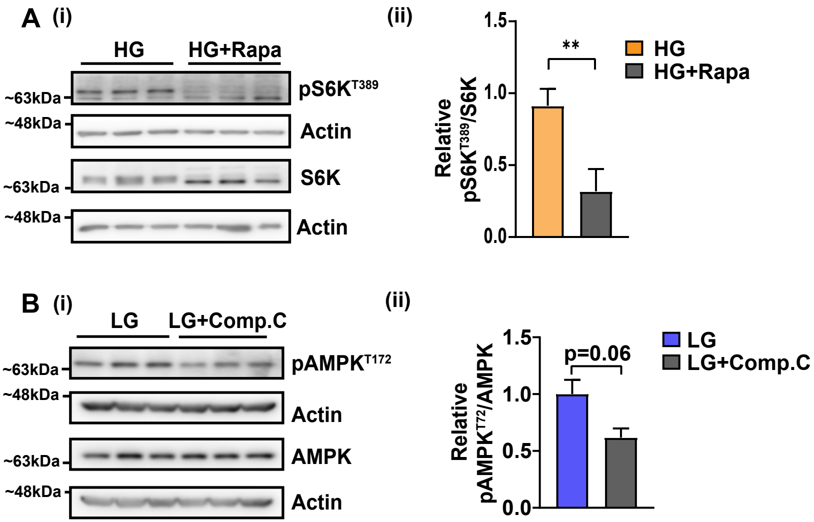

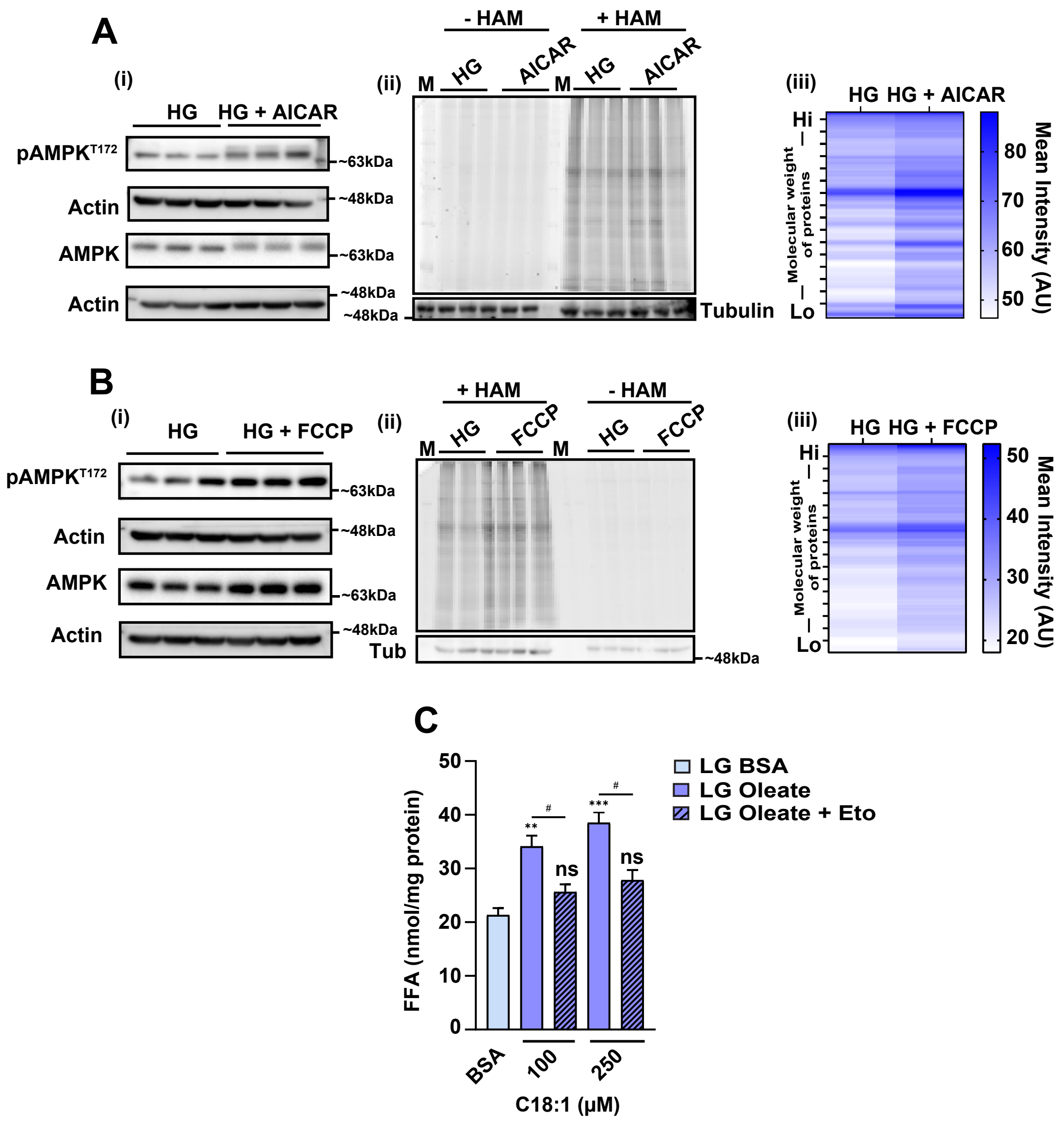

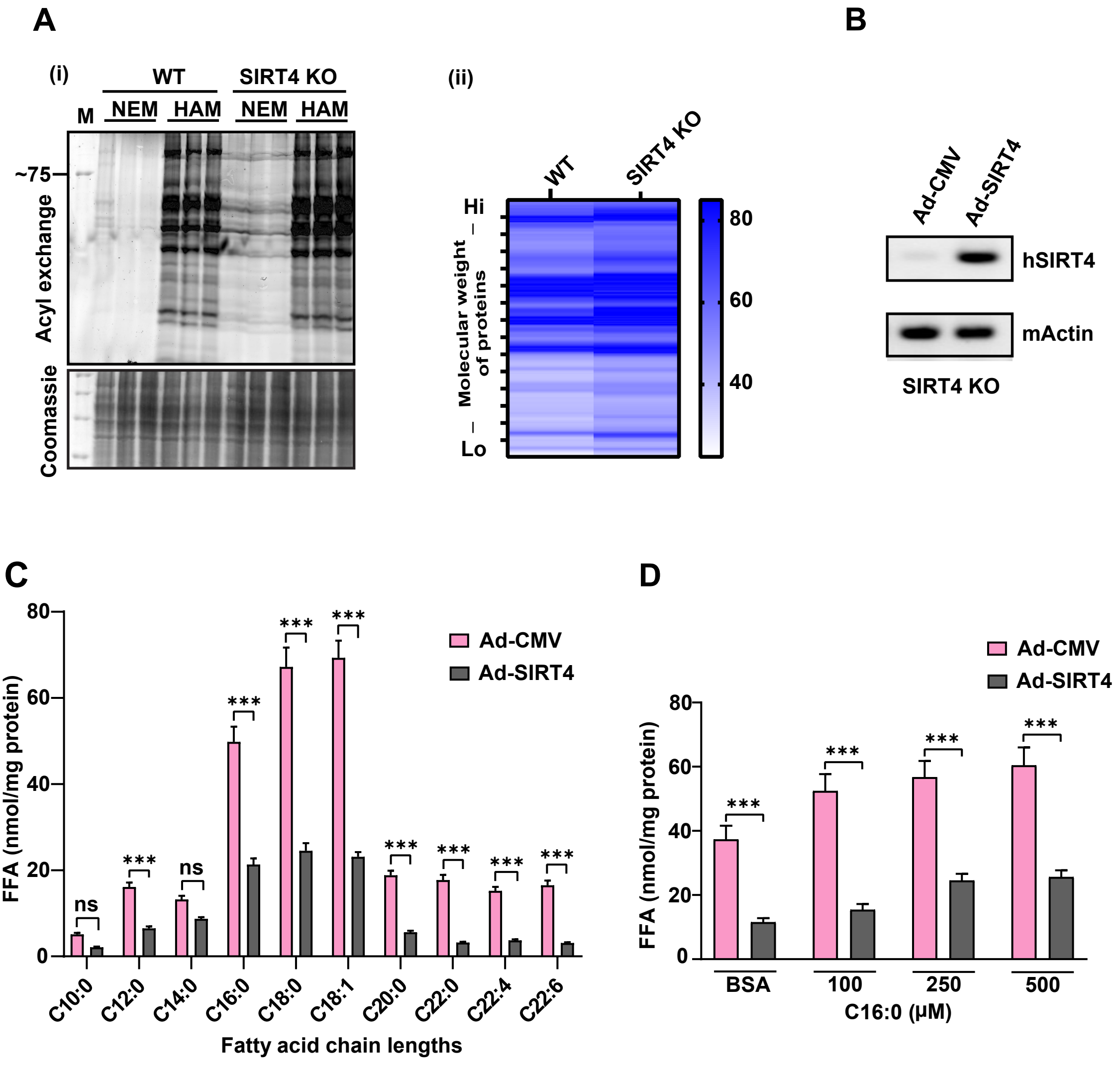

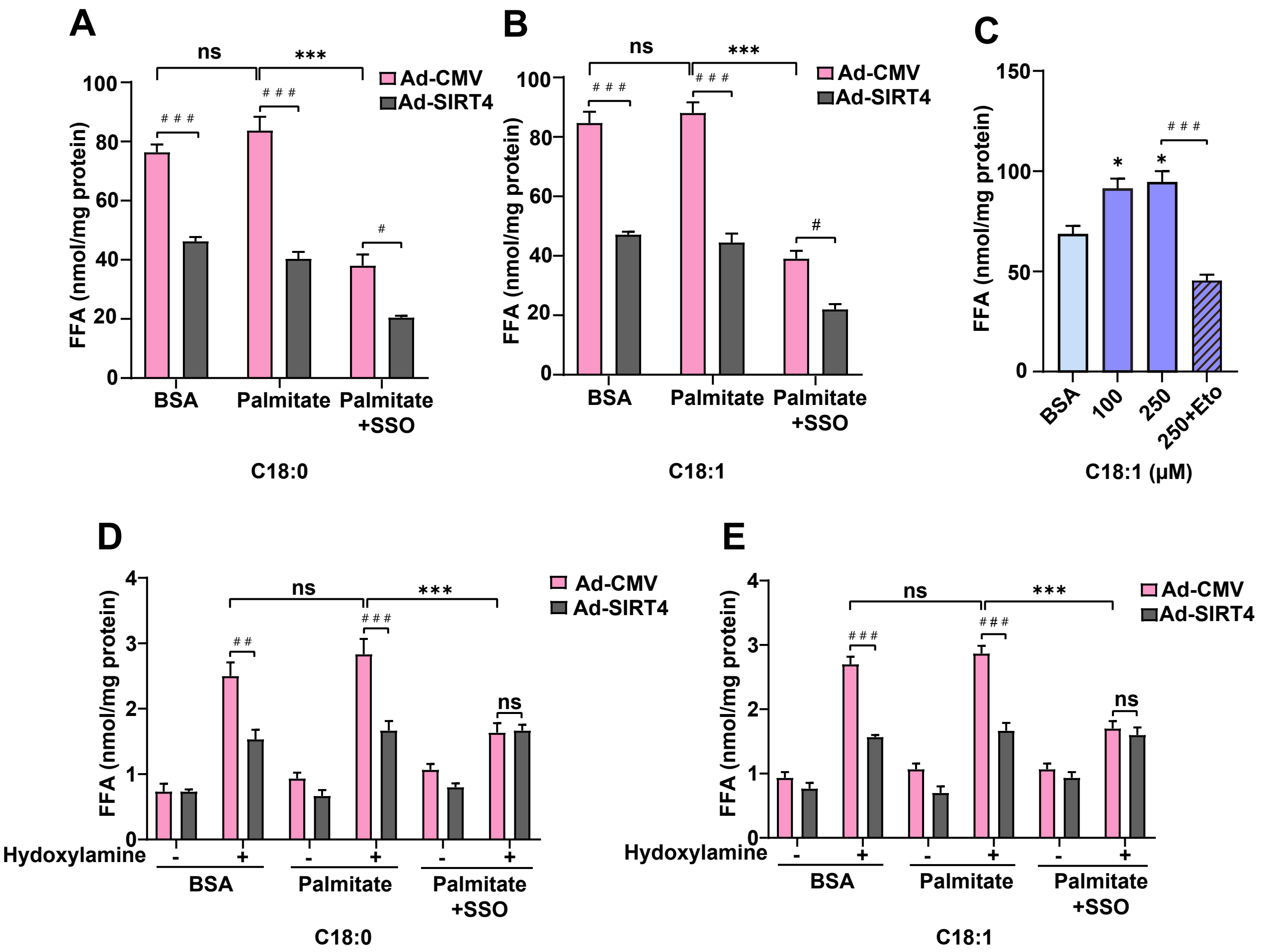
